# Choice of fluorophore affects dynamic DNA nanostructures

**DOI:** 10.1101/2020.12.12.422444

**Authors:** Kevin Jahnke, Helmut Grubmüller, Maxim Igaev, Kerstin Göpfrich

## Abstract

The ability to dynamically remodel DNA origami structures or functional nanodevices is highly desired in the field of DNA nanotechnology. Concomitantly, the use of fluorophores to track and validate the dynamics of such DNA-based architectures is commonplace and often unavoidable. It is therefore crucial to be aware of the side effects of popular fluorophores, which are often exchanged without considering the potential impact on the system. Here, we show that the choice of fluorophore can strongly affect the reconfiguration of DNA nanostructures. To this end, we encapsulate a triple-stranded DNA (tsDNA) into water-in-oil compartments and functionalize their periphery with a single-stranded DNA handle (ssDNA). Thus, the tsDNA can bind and unbind from the periphery by reversible opening of the triplex and subsequent strand displacement. Using a combination of experiments, molecular dynamics (MD) simulations, and reaction-diffusion modeling, we demonstrate for twelve different fluorophore combinations that it is possible to alter or even inhibit the DNA nanostructure formation – without changing the DNA sequence. Besides its immediate importance for the design of pH-responsive switches and fluorophore labelling, our work presents a strategy to precisely tune the energy landscape of dynamic DNA nanodevices.

## Introduction

DNA nanotechnology has been highly successful in repurposing the iconic DNA double helix to create programmable molecular architectures. Once focused on static shapes, dynamic and stimuli-responsive DNA nanoscale devices are gaining a large surge of interest for various applications^[1]^ – from sensors,^[2–4]^ biocomputing algorithms,^[5]^ and drug delivery systems^[6,7]^ to programmable robotic modules^[8,9]^ and functional components for synthetic cells. ^[10–12]^ In a vast majority of such reconfigurable systems, dynamics is achieved using strand displacement reactions,^[13,14]^ flexible single-stranded hinges,^[15]^ or stimuli-responsible DNA modifications ^[16,17]^ and sequence motifs.^[4,18]^ The ability to reversibly actuate artificial structures at the nanoscale is therefore at the core of dynamic DNA nanotechnology. Direct measurements of conformational changes in aqueous solutions are often conducted with Forster resonance energy transfer (FRET) or fluorescence microscopy. ^[19–21]^ These methods can provide a readout of the overall conformational state of the structure, for example, open *versus* closed, or bound *versus* unbound. Hence, the use of fluorescent dyes is commonplace to validate and quantify the functionality of the DNA-based devices. Fluorophore-tagged DNA nanostructures have also been used as nanoscopic rulers for fluorescence microscopy^[22]^ and to enable the acquisition of super-resolution images with DNA-PAINT.^[23]^ Factors like solubility, photostability and excitation/emission spectra usually play the decisive role in choosing the suitable dye, while potential side effects on the DNA conformation such as overstabilization of DNA duplexes^[24]^ or specific fluorophore-DNA interactions^[25]^ are not the main concern.

Here, we show that the choice of the fluorophore itself can alter the equilibrium conformation and even inhibit a desired dynamic response. We use a pH-responsive triple-stranded DNA motif (tsDNA) combined with a strand-displacement reaction to exemplify that the dynamics can be strongly influenced by the choice of the fluorophore. With all-atom molecular dynamics (MD) simulations, we show that fluorophore-dependent conformational dynamics of the single-stranded DNA (ssDNA) contributes to this observation. By releasing caged protons inside droplet-based confinement, we find that also the duplex dissociation is affected by the fluorophore. Using a reaction-diffusion model, we derive the apparent dissociation constant for 12 different experimentally tested fluorophore combinations. A profound knowledge about the effect that fluorophores and other chemical modifications have on the dynamics of a DNA-based system can be leveraged to realize the desired functionality.

## Materials and Methods

### DNA sequence design

The DNA sequences were adapted from Green et al.^[4]^ To enable self-assembly at the droplet-periphery, the ssDNA (termed ‘Regulator’ in Green et al.) was modified with a cholesterol-tag (sequence: 5’ Cy3/Alexa488/Cy5/-ACCAGACAATACCACACAATTTT-CholTEG 3’, HPLC purified). The tsDNA (termed ‘Sensor’ in Green et al.) contained the triple-stranded DNA motif as well as a stem loop complementary to the ssDNA. A fluorophore modification was added to its 5’ end (sequence: 5’ Cy5/Cy3/Atto488/Atto647-TTCTCTTCTCGTTTGCTCTTCTCTTGTGTGGTATTGTCTAAGAGAAGAG 3’, HPLC purified). Both DNA sequences were purchased from Biomers or Integrated DNA Technologies and dissolved in ultrapure water (Milli-Q) to exclude the impact of DNA storage buffer on the pH.

### Formation of DNA-containing water-in-oil droplets

For the formation of water-in-oil droplets, the DNA-containing aqueous phase was layered on top of the oil phase in a volumetric ratio of 1:3 within a microtube (Eppendorf). Droplet formation was induced by manual shaking for about 4 s as described earlier.^[26]^ For the oil-phase, 1.4 vol% of perflouro-polyether-polyethylene glycol (PFPE-PEG) block-copolymer fluorosurfactants (008-PEG-based fluorosurfactant, Ran Biotechnologies, Inc.) dissolved in HFE-7500 oil (DuPont) was used. The interfacially active surfactants stabilize the droplets. The aqueous phase was composed of 10 mM MgCl_2_ and 250 mM potassium phosphate buffer adjusted to pH values from 4.3 to 8.0. Cholesterol-tagged ssDNA and the tsDNA were added to the aqueous phase at concentrations of 1.66μM and 1.25 μM, respectively, if not stated otherwise. ssDNA was added in excess to ensure that there are sufficient binding sites for the tsDNA. Other contents were encapsulated by adding them to the aqueous phase as described in text.

### Confocal fluorescence microscopy

For confocal microscopy, the DNA-containing droplets were sealed in a custom-built observation chamber and imaged 10 min after encapsulation using a confocal laser scanning microscope LSM 880 or LSM 800 (Carl Zeiss AG). The pinhole aperture was set to one Airy Unit and experiments were performed at room temperature. The images were acquired using a 20x objective (Plan-Apochromat 20x/0.8 M27, Carl Zeiss AG). Images were analyzed and processed with ImageJ (NIH, brightness and contrast adjusted).

### Light-triggered proton release

To dynamically decrease the pH inside individual compartments, we co-encapsulated 40 mM NPE-caged sulfate (Santa Cruz Biotechnology), which undergoes photolysis upon illumination with light of the wavelength 405 nm and releases a proton. For the investigation of the detachment kinetics, 2μM ssDNA and 1.5 μM tsDNA were mixed with 20mM potassium phosphate buffer at pH 8 and 5 mM MgCl2. The use of the buffer ensures the same starting conditions and delays the acidification, which facilitates the imaging and analysis of the tsDNA fluorescence. After encapsulation, a subset of droplets was illuminated with 20% of the power of a 5 mW 405 nm laser diode while simultaneously recording the detachment of the Cy5-labelled tsDNA. The field of view, the laser intensity and all additional imaging conditions were kept the same.

### Image analysis

The tsDNA fluorescence inside the droplet and at the droplet periphery was analysed with a custom-written ImageJ macro. Droplets were identified and assigned a circular region of interest from which the droplet radius *r*_0_ was calculated. The intensity within the droplet center, *I*_in_, was defined as the mean intensity within a circle of radius *r*_in_ = 0.5*r*_0_. The intensity at the droplet periphery *I*_peri_ was quantified by measuring the maximum intensity along a line orthogonal to the droplet circumference. This analysis was repeated 20 times every 18° along the droplet circumference, and the mean value taken as *I*_peri_. Following the determination of the droplet intensities I_in_, they were plotted with Prism 8 (GraphPad) and fitted using a sigmoidal function of the form: *I*_in_ = *I*_min_ + (*I*_min_ – *I*_max_)/(1 + 10^−*α*(*x*_turn_−*x*)^), with *α* being the decay constant and *x*_turn_ the turning point of the fit.

### Atomistic simulations of unlabeled ssDNA

To provide a realistic description of ssDNA dynamics both in the presence and in the absence of fluorescent dyes, we first performed a series of simulations for the dye-free ssDNA using the Parmbsc1 flavour^[27]^ of the standard Amber 99SB force field^[28]^ with CUFIX nonbonded corrections^[29]^ and ion parameter corrections by Joung and Cheatham.^[30]^ We also used TIP3P as the water model in our simulations.^[31]^ The simulations were initiated from single-stranded helical structures built with Chimera (v. 1.14).^[32]^ The starting structures were solvated in TIP3P water in a dodecahedron box with an edge length of 9.0 nm, yielding a system with approximately 50 000 atoms. Ion concentrations were set to 250 mM NaCl and 10 mM MgCl_2_ to mimic the experimental conditions.

Subsequent MD simulations were performed with GROMACS 2019.6.^[33]^ Lennard-Jones and short-range electrostatic interactions were calculated with a 1.0-nm cutoff, while long-range electrostatics was treated using particle-mesh Ewald summation^[34]^ with a 0.12-nm grid spacing. Hydrogen bond lengths were constrained using the LINCS algorithm.^[35]^ Velocity rescaling^[36]^ with a heat bath coupling constant of 1.0 ps was used to control the temperature for solute and solvent separately. Center-of-mass correction was applied to solute and solvent separately every 100 steps. Energy minimization was followed by a short equilibration for 1 ns in the NVT ensemble (T = 100 K) and with position restraints applied to the solute’s heavy atoms and a 1-fs integration time step. Next, the temperature was increased to T = 300 K, and the system was equilibrated for 5 ns (2-fs time step), while keeping the pressure at 1 atm using the Berendsen barostat^[37]^ with a 1-ps coupling constant. The position restraints were then slowly released during 20 ns of equilibration in the NPT ensemble (T = 300 K, p = 1 atm, 2-fs time step) using the Parrinello-Rahman barostat.^[38]^ This initial equilibration step was followed by a total of 17 independent production runs, each being 5 *μ*s long. The first ~1 μs of the trajectories were discarded to exclude the initial relaxation towards the equilibrium state. Unless specified differently, all trajectory analyses were performed with Python (v. 2.7 available at https://www.python.org/), VMD (v. 1.9.2)^[39]^ and Chimera (v. 1.14).^[32]^

### ssDNA simulations with fluorescent dyes covalently attached

Parameters and energy-minimized structures for common Alexa and Cy fluorescent dye families were derived from the AMBER-DYES library^[40]^ that is compatible with most Amber force fields. Alexa488, Cy3 (water-soluble) and C5 (water-soluble) dyes were attached to the 5’ end of the ssDNA via a neutral lysine linker. To this end, the capping H5T atom of the 5’ nucleotide was removed, and the C atom of the linker backbone was bonded with the O5’ atom of the 5’ nucleotide. Since in the Amber formalism, the 5’ and 3’ nucleotides possess non-integer charges (−0.3 e and −0.7 e, respectively; unlike the regular nucleotides that have a charge of −1.0 e), the resulting dye-ssDNA construct had a slightly non-integer charge. To account for this, the residual small charge was redistributed over the O5’, C5’, C4’, C3’, O4’, C1’, and C2’ atoms of the 5’ nucleotide (sugar backbone).

To accommodate the larger dye-ssDNA, the size of the simulation box was increased to 12 nm, yielding a system with about 120 000 atoms. All subsequent simulations were done under the same conditions as for the unlabeled ssDNA. For the dye-free simulations, multiple 6 μs production runs were performed and the first ~1 μs were discarded as equilibration time. A summary of the simulated systems is given in Table 1.

**Table 1:** Summary of dye-free and dye-labeled ssDNA simulations.

| Force field | System | Duration |
| --- | --- | --- |
| Parmbsc1 + TIP3P | no dye | $17 \times 5 \mu s$ |
| Parmbsc1 + TIP3P | Cy3 | $6 \times 6 \mu s$ |
| Parmbsc1 + TIP3P | Cy5 | $6 \times 6 \mu s$ |
| Parmbsc1 + TIP3P | Alexa 488 | $6 \times 6 \mu s$ |

### Determination of the apparent dissociation constant

The equilibrium distribution of ssDNA and tsDNA molecules in a droplet can be described mathematically using a reaction-diffusion system of equations in which the binding sites (ssDNA attached to the droplet periphery), and hence also the binding and dissociation reactions, are localized in a narrow volumetric layer near the spherical droplet surface. ^[41,42]^ Briefly, if *S*_tot_ and *T*_tot_ are the total concentrations of ssDNA and tsDNA in the droplet, respectively, *T*_eq_ is the steady-state concentration of tsDNA in equilibrium, and *K*_D_ is the dissociation constant defining the ssDNA/tsDNA binding equilibrium, the ratio between the peripheral and inner intensity of tsDNA fluorescence can be expressed as:

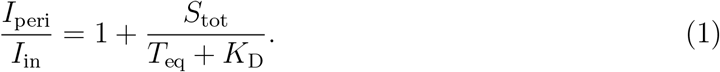

Here, both *I*_peri_ and *I*_in_ are per-area intensities averaged over 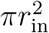 and 2*πr*_0_*ε*, respectively, where *ε* is the apparent thickness of the reaction layer (determined from confocal images as described in the Supplementary Text S1). The steady-state concentration *T*_eq_,

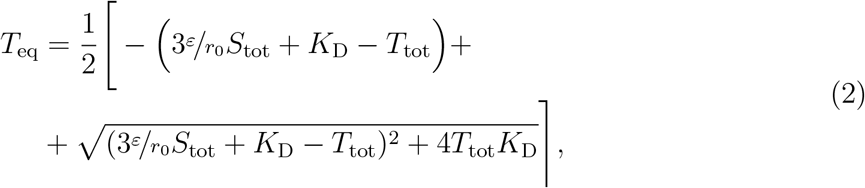

is obtained by simultaneously requiring that the ssDNA/tsDNA binding has attained equilibrium and that the number of tsDNA molecules in the droplet is conserved. The apparent *K*_D_ values presented in Figure 4 were determined using Eqs. 1 and 2, and the corresponding errors were calculated using basic error propagation rules and measured uncertainties of *I*_peri_, *I*_in_, *r*_0_, and *ε*. A detailed mathematical description of the model, estimation of *ε,* and error analysis are given in the Supplementary Texts S1 and S2.

### Radius of gyration distributions and estimations of confidence intervals

The gyration radii (*R_g_*) of ssDNA were calculated using the gmx gyrate tool included in the GROMACS package. The probability distributions *p*(*R_g_*) shown in Figure 2 were then computed by binning the corresponding data sets and normalizing the histograms. Confidence intervals for *p*(*R_g_*) were estimated using bootstrap analysis.^[43]^ To this end, we used the obtained distributions to bootstrap 10^6^ new random *R_g_* samples (each consisting of 10^5^ data points) such that the newly generated data is distributed according to *p*(*R_g_*) and properly correlated with the autocorrelation time estimated from the original *R_g_* trajectories.

## Results

### Fluorophore modification influences pH response

We set out to test the impact of fluorophores on the dynamics of DNA nanostructures. For this purpose, we employed a popular triple-stranded DNA motif (tsDNA)^[4]^ as an example. Its reversible pH-responsive actuation works as follows: At basic pH, the Hoogsten-interactions which stabilize the triple-stranded configuration are weaker than at acidic pH. Therefore, an increase in pH leads to unwrapping of the third strand which previously stabilized the duplex. This, in turn, lowers the energy barrier for a strand displacement reaction with a single-stranded DNA (ssDNA), which was designed to be complementary to the hairpin loop of the tsDNA. Thus, a stable DNA duplex can form between the tsDNA and the ssDNA (Figure 1**A**).^[4]^ To monitor this process, we modified the ssDNA with a cholesterol-tag and encapsulated it together with the tsDNA into water-in-oil droplets. Upon encapsulation, the ssDNA self-assembled at the droplet periphery due to hydrophobic interactions between the cholesterol-tag and the droplet-stabilizing surfactants.^[11]^ Thereby, we obtained an attachment handle, which reversibly recruits the tsDNA to the periphery at basic pH (Figure 1**B**). In contrast to Förster Resonance Energy Transfer (FRET), which is commonly employed to monitor the pH dynamics,^[21]^ our system provides freedom regarding the choice of fluorophores - which is absolutely necessary for us to study the impact of different fluorophore combinations. We directly visualized tsDNA binding and investigated the impact of fluorophore modifications on the pH dynamics. At the same time, this system provides a strategic blueprint for the pH-sensitive recruitment of components to the membrane - an interesting function in itself, in particular concerning the bottom-up construction of synthetic cells.^[44]^

**Figure 1:**
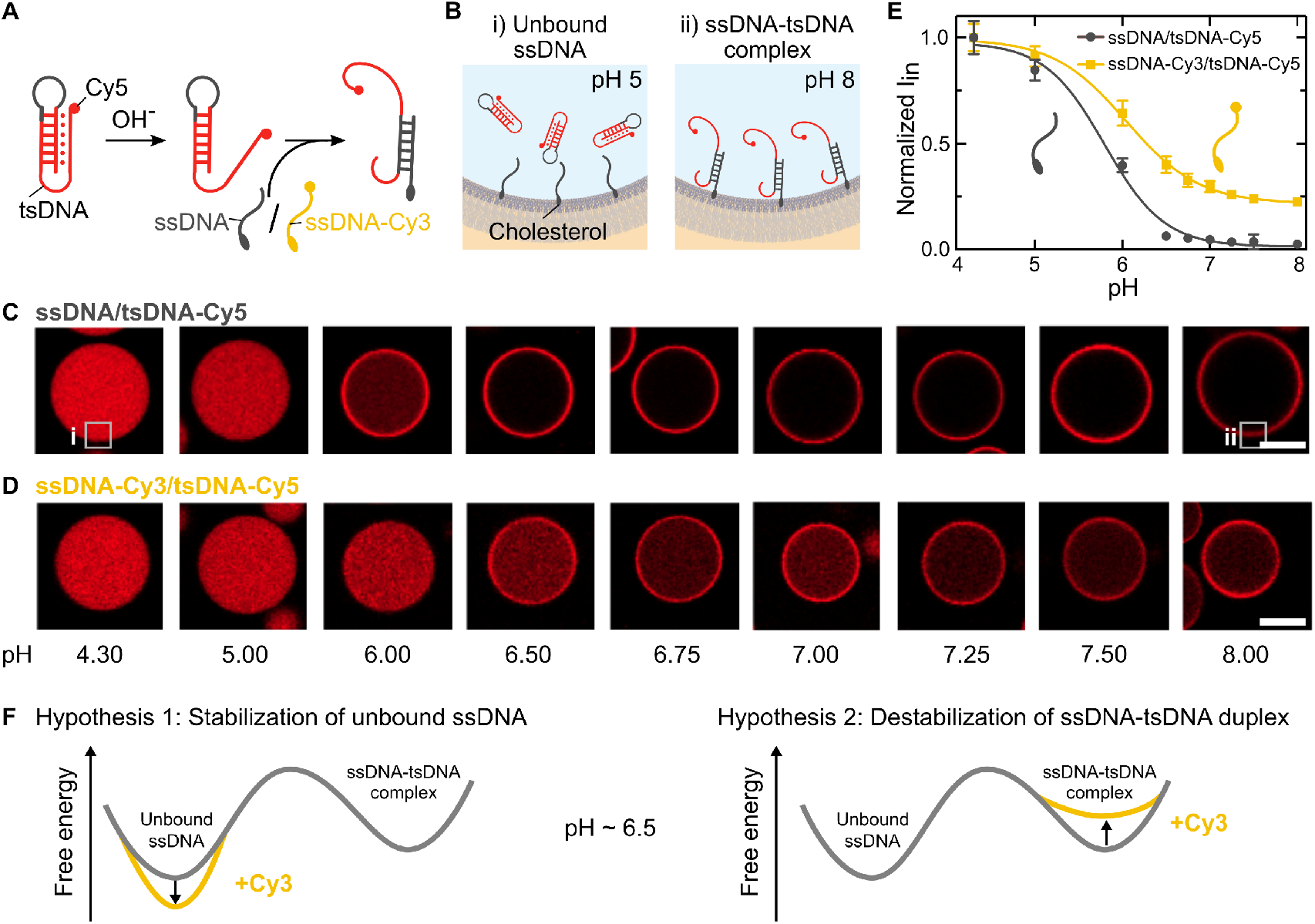
Fluorophore modification influences pH response. **A** Sketch of the pH responsive DNA motif. A Cy5-labeled triple-stranded DNA (tsDNA) opens up at basic pH, lowering the energy barrier for strand displacement and hence for complementary base pairing with a cholesterol-tagged single-stranded DNA (ssDNA). **B** This process can be monitored in water-in-oil droplets. The cholesterol-tagged ssDNA self-assembles at the droplet periphery, whereas cholesterol-free tsDNA remains homogeneously distributed within the droplet at acidic pH and attaches to the droplet periphery at higher pH values. **C, D** Representative confocal images of water-in-oil droplets containing Cy5-labeled tsDNA (red, *λ_ex_* = 647 nm) and unlabeled ssDNA (**D**) or Cy3-labeled ssDNA (**E**) at different pH values. Attachment of the tsDNA is shifted to higher pH values if the ssDNA is labelled with Cy3. Scale bars: 20 μm. **E** Normalized fluorescence intensity of the Cy5-labeled tsDNA inside the droplet (periphery excluded) at different pH values for unlabeled (gray) and Cy3-labeled ssDNA (yellow). Error bars correspond to the standard deviation of the intensities of n ≥ 9 droplets. Solid lines represent sigmoidal fits revealing a turning point at pH 5.80 ± 0.09 and 6.05 ± 0.04, respectively. **F** Free energy profile illustrating potential hypotheses for the altered behaviour of the Cy3-tagged ssDNA compared to the unlabelled ssDNA.

**Figure 2:**
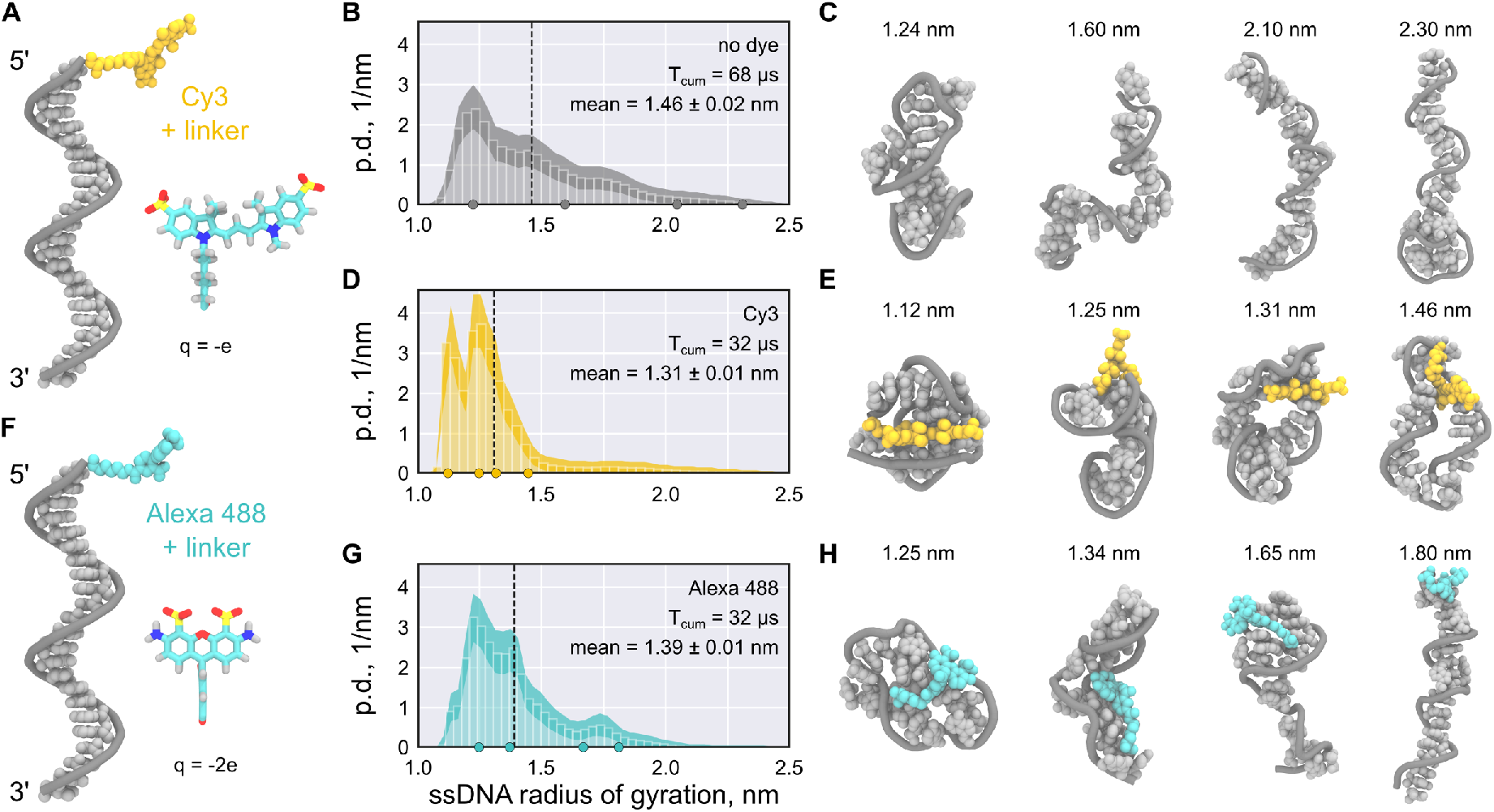
MD simulations suggest that fluorophore labeling can stabilize more compact ssDNA conformations. **A, F** Starting conformation of Cy3-(A, yellow) and Alexa488-labeled (F, turquoise) ssDNA (gray). The chemical structure of the fluorophore and its net charge are shown as an inset. **B, D, G** Probability density (p.d.) distributions of the gyration radius of ssDNA without dye (B), labeled with Cy3 (D), and labeled with Alexa488 (G). The shaded areas indicate the 95% confidence intervals estimated using bootstrapping (see Methods). The black dashed lines indicate the means of the distributions, T_cum_ the cumulative simulation time. **C, E, H** Representative structure snapshots of the unlabelled ssDNA (C), the Cy3-ssDNA (E) and the Alexa488-ssDNA (H). Positions of the selected snapshots within the corresponding distributions are marked with dots in the probability density distributions.

Confocal imaging revealed that attachment of the tsDNA to the compartment periphery is shifted to higher pH values if the ssDNA carries a Cy3 compared to the unlabelled ssDNA (Figure 1**C,D**). The images show the equilibrated state (Figure S1) and we confirmed that the shift is not due to interactions of the Cy3 with the surfactant layer (Figure S2). We quantified this effect by extracting the normalized intensity inside the droplets (I_in_, periphery excluded) from the confocal images (Figure 1**E**). Importantly, we could reproduce the sigmoidal pH response curve that was reported for FRET-based detection.^[45]^ The turningpoint of the pH-sensitive ssDNA-tsDNA binding curve for unlabelled ssDNA was around 5.80 ± 0.09, whereas it shifted significantly to 6.05 ± 0.04 for the Cy3-modified ssDNA (2.53*σ*). Even at pH 8 not all tsDNA was bound to the droplet periphery for the Cy3-modified ssDNA.

While it is well known that the pH turning point can be shifted by changing the DNA sequence,^[45]^ it was not known that the same can be achieved by changing the fluorophore modification alone. This striking observation can be explained by either of the two following hypotheses as illustrated in Figure 1**F**: 1) A fluorophore modification on the ssDNA causes overstabilization of the free ssDNA state by making its equilibrium ensemble more compact and, therefore, less accessible for base paring. 2) The fluorophore modification destabilizes the ssDNA-tsDNA complex, thereby raising the bound state in free energy (relative to the unbound one). First, we tested Hypothesis 1 with all-atom molecular dynamics (MD) simulations, subsequently we examined Hypothesis 2 with experiments.

### MD simulations reveal reduction of ssDNA accessibility by fluorophore modification

To test Hypothesis 1, we used all-atom MD simulations to probe the secondary structure of the Cy3-labelled ssDNA (Figure 2**A**) and compared it to the unlabelled ssDNA. First of all, the unlabelled ssDNA yielded a very broad probability density distributions for the radius of gyration (Figure 2**B**), which is a direct measure of the DNA’s compactness. The distributions for the unlabelled ssDNA show a significant fraction of extended structures, in which the DNA bases are accessible for complementary base pairing (see also representative snapshots in Figure 2**C** and Video S1). On the contrary, the Cy3-labelled ssDNA (Figure 2**A**) yielded a distinctively different probability density distribution for the radius of gyration (Figure 2**D**), which reflects a much lower propensity for extended conformations. The bases of the Cy3-labelled ssDNA were found to be wrapped around the fluorophore, most likely due to stacking interactions between the ssDNA bases and the aromatic groups of Cy3 (Video S2). This entangled conformation renders the ssDNA less accessible for complementary base pairing. An overstabilization of the unbound ssDNA means a lower free energy of the ssDNA compared to the ssDNA-tsDNA complex. This would explain our experimental observations in line with Hypothesis 1. Note that Cy5-labelled ssDNA favored similarly compact conformations wrapped around the dye, which further indicates that the aromatic groups of Cy dyes tend to interact with ssDNA base pairs (Figure S3, Video S3).

To test if weaker dye-ssDNA interactions would restore expanded conformations of the ssDNA in our simulations, we used an Alexa 488 dye. We selected an Alexa dye (Figure 2**F**), because its chemical structure is considerably different compared to Cy3, which may render it less prone to base stacking interactions. Moreover, Alexa dyes are more hydrophilic due to their two negative charges. We found that the mean radius of gyration for an Alexa488-modified ssDNA laid between that of the unmodified and the Cy3-modified ssDNA (Figure 2**G**). The MD snapshots show that the fully extended conformation, where the bases are accessible, was partially recovered (Figure 2**H**, Video S4), improving the accessibility of the strand for complementary base pairing.

Taken together, our simulations suggest that a single fluorophore modification on ssDNA can significantly change the DNA’s conformation. The more compact conformation of dye-labeled ssDNA effectively increases the free energy cost for expansion required for duplex formation with tsDNA. Thus, our simulations support Hypothesis 1. Importantly, the ssDNA sequence is random such that the observations can likely be generalized for a broad spectrum of DNA sequences.

### Dissociation kinetics show fluorophore dependence

As a next step, we investigated the duplex dissociation process to see if the fluorophore modification affects the dissociation constant after duplex formation (Hypothesis 2). Since all-atom MD simulations cannot describe this reaction due to the limited timescales, we studied the detachment of the tsDNA from the compartment periphery experimentally. We implemented an approach where we achieved light-triggered release of protons in individual compartments – providing full spatio-temporal control over the acidification process. For this purpose, we used NPE-caged-sulfate, which breaks up into a sulfate and a proton upon photolysis.^[46]^ To prove that NPE-caged sulfate can be used to decrease the pH inside the droplets, we first encapsulated it together with the pH-sensitive dye pyranine at pH 8 and locally illuminated individual droplets with a 405 nm laser (Figure 3**A**). The pyranine emission upon 488 nm excitation decreased, confirming the successful pH decrease inside the droplets from initially pH 8 to under pH 5. The buffer kept the pH constant until its capacity is exceeded after approximately 20 s. Then, the pH decreased until most of the NPE-caged sulfate underwent photolysis and hence the pH approached a constant value after ~50s. We used this dynamic light-mediated acidification mechanism to detach the tsDNA from the droplet periphery. At t=0 s, the tsDNA was bound to the ssDNA at the droplet periphery (Figure 3**B**) and completely detached within 50 s of illumination. Upon detachment, the triplex conformation of the tsDNA was restored. In order to assess the detachment kinetics, we monitored the normalized tsDNA-Cy5 intensity for unmodified, Cy3-modified and Alexa488-modified ssDNA inside the droplet over time (Figure 3**C**).

**Figure 3:**
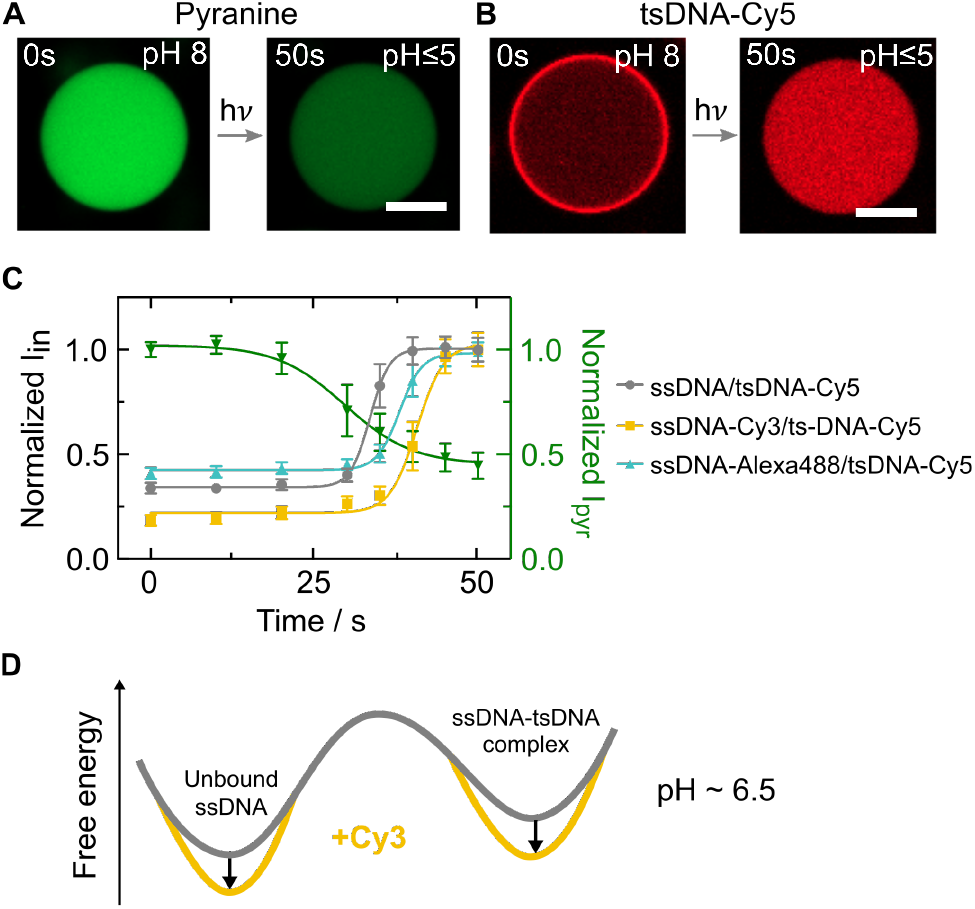
Light-mediated acidification of water-in-oil droplets reveals fluorophore-dependent duplex dissociation kinetics. **A** Confocal images of the pH-sensitive dye pyranine (50 μM, *λ_ex_* = 488 nm, not coupled to DNA) encapsulated into water-in-oil droplets at pH 8. Light-triggered uncaging of NPE-caged sulfate (λ_*ex*_ = 405 nm) leads to proton release causing a rapid pH drop from 8 to under 5 within 50 s. The pH drop can be monitored as a decrease in pyranine fluorescence. **B** Representative confocal images of Cy5-labeled tsDNA (λ_*ex*_ = 647 nm) encapsulated together with cholesterol-tagged ssDNA into water-in-oil droplets at pH 8. During acidification, the tsDNA detaches from the droplet periphery as the triplex state is energetically favoured. Scale bars: 20 μm. **C** Normalized fluorescence intensity of the tsDNA inside the droplet (periphery excluded) over time for unlabeled, Cy3-labeled and Alexa488-labeled ssDNA as well as pyranine (right axis). Error bars correspond to the standard deviation of the intensities of n ? 23 droplets for the DNA experiments and n = 6 droplets for pyranine experiments. Solid lines represent sigmoidal fits with turning points at 33.5s ± 0.1s (unmodified ssDNA), 40.7s ± 0.5s (ssDNA-Cy3) and 38.0s ± 0.3s (ssDNA-Alexa488). Note that the decay times *t_d_* = 1/α are similar for all fluorophores 345s ± 0.24 s (unmodified ssDNA), 4.76s ± 1.13s (ssDNA-Cy3) and 4.00s ± 0.48s (ssDNA-Alexa488). **D** Free energy profile illustrating our conclusion that both equilibrium states are stabilized by the presence of a dye on the ssDNA.

**Figure 4:**
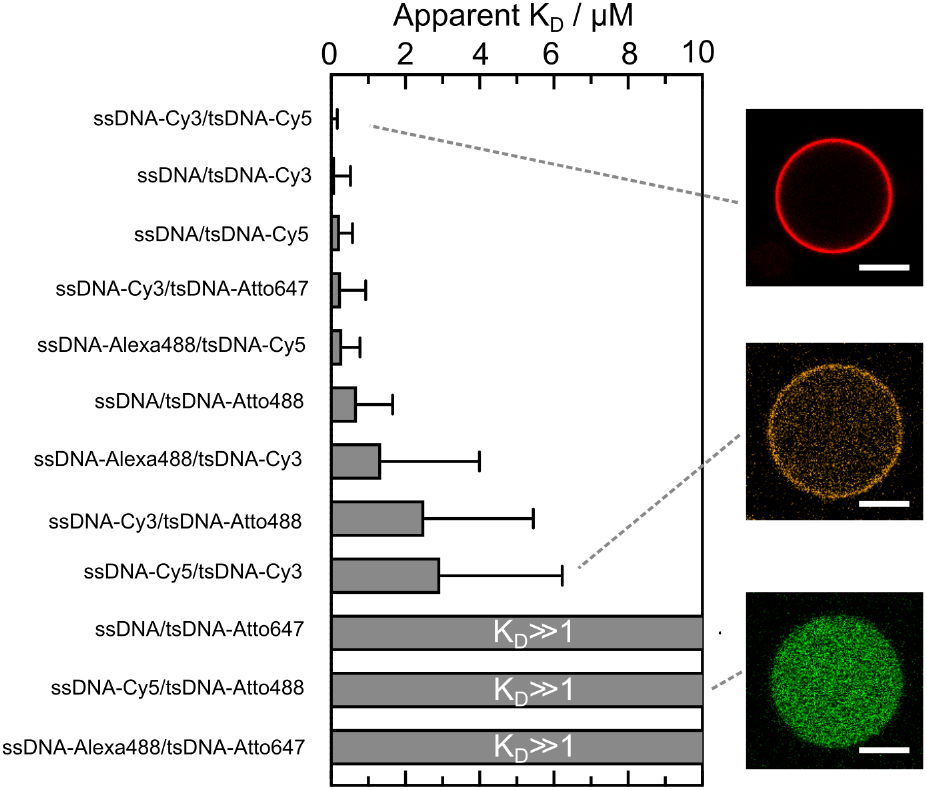
Histogram depicting the apparent dissociation constants K_D_ for 12 different ssDNA/tsDNA combinations at pH 8 with variable fluorophore modifications including Alexa488 (λ_*ex*_ = 488nm), Atto488 (λ_*ex*_ = 488nm), Cy3 (λ_*ex*_ = 561 nm), Cy5 (λ_*ex*_ = 647nm) and Atto647 (λ_*ex*_ = 647nm). Error bars correspond to the standard deviation of n = 1173 evaluated droplets. Confocal images of three fluorophore combinations depicting strong (ssDNA-Cy3/tsDNA-Cy5), intermediate (ssDNA-Cy5/tsDNA-Cy3) and almost no binding to the droplet periphery (ssDNA-Cy5/tsDNA-Atto488). The apparent K_D_ is strongly influenced by fluorophore modifications on ssDNA and tsDNA up to the point of almost full inhibition of binding, which results in K_D_ ≫ 1.

Following proton-release, the tsDNA detached from the ssDNA for all tested fluorophore modifications (Video S5).

The decay times *t_d_* = 1/*α* of the sigmoidal fits were comparable for all three fluorophore modifications, indicating similar detachment kinetics. However, detachment (i.e. duplex dissociation) occurred at different time points, hence at different pH values - again pointing towards an altered binding equilibrium. Detachment from the unlabelled ssDNA happened earlier (i.e. at higher pH) indicating that a fluorophore label is stabilizing the ssDNA-tsDNA complex.

Taken together, the results obtained so far suggest that fluorophore modifications, in particular Cy-dyes, stabilize not only the unbound ssDNA (Hypothesis 1) but also the ssDNA-tsDNA duplex as illustrated in the free energy profile in Figure 3**D**. However, the stabilization of compact ssDNA conformations is likely stronger, which explains the observed shift of the pH switching point. This is effectively increasing the energy barrier for the dynamic switching of fluorophore-labelled DNA.

### Reaction-diffusion modelling reveals impact of fluorophores on apparent dissociation constant

Finally, having shown that a fluorophore modification on the ssDNA has a significant influence on the pH switching point, we now additionally tested the impact of fluorophore modifications on tsDNA. For this purpose, we investigated twelve different fluorophore combinations on ssDNA and tsDNA. To quantitatively compare the impact of different fluorophores, we developed an analytical model to derive the apparent dissociation equilibrium constant *K*_D_ = *k*_off_/*k*_on_ at a fixed pH for each individual fluorophore combination. For this purpose, we derived a reaction-diffusion model for spherical compartments (Text S2). It allowed us to determine the apparent dissociation constant *K*_D_ by extracting the droplet radius, the peripheral and the inner intensity of the tsDNA from confocal images with known total concentrations of DNA. We tested combinations of five different fluorophores, namely Cy3, Cy5, Alexa488, Atto488 and Atto647 as well as unlabeled ssDNA on the apparent *K*_D_ (Figure 4). Note that the use of an unlabelled tsDNA was not possible because it would inhibit the monitoring with confocal microscopy.

Remarkably, *K*_D_ varied dramatically for the different combinations. Most striking was the fact that binding is almost fully inhibited for certain fluorophore combinations, like ssDNA/tsDNA-Atto647, ssDNA-Cy5/tsDNA-Atto488 and ssDNA-Alexa488/tsDNA-Atto647 with *K*_D_ ≫ 1. On the other hand combinations like ssDNA-Cy3/ts-DNACy5, ssDNA/tsDNA-Cy3 and ssDNA/tsDNA-Cy5 bound very efficiently as expected at pH 8. As a general trend, we deduce that Cy-dyes on the tsDNA seemed to lead to a lower apparent *K*_D_ compared to Atto-dyes. Furthermore, it is surprising that the permutation of two Cy-dyes on ssDNA and tsDNA lead to a different apparent *K*_D_. While ssDNA-Cy3/tsDNA-Cy5 attached very efficiently, we obtained intermediate *K*_D_’s for ssDNA-Cy5/tsDNA-Cy3. Confirming our observations, the permutation of the tsDNA fluorophore influenced the pH hysteresis (Figure S4) and the dynamic detachment in experiments using NPE-caged sulfate (Figure S5).

Taking all these observations into account, we propose that not only a fluorophore modification on the ssDNA but also on the tsDNA affects the dynamics of pH-responsive DNA nanostructures up to a point that binding is inhibited. The choice of fluorescent dyes can thus be exploited to shape the energy landscape for dynamic DNA nanostructures and to shift the equilibrium towards the bound or the unbound state.

## Discussion

One of the most exciting tasks in the field of DNA nanotechnology is the construction of dynamic molecular devices that can perform mechanical motion upon stimulation. The foundation for this work is an experimental readout, which is suitable to track dynamic reconfiguration in space and time. Fluorescence microscopy techniques, such as superresolution imaging or FRET, are ideally suited for *in situ* measurements on active DNA origami structures. The fluorophore is normally selected to match the optical setup rather than the DNA nanostructure itself and exchanged as required by the experiment. Here, we determined why the exchange of fluorophores on dynamic DNA nanostructures can lead to a considerably different experimental outcome. We used a popular pH-sensitive DNA motif combined with a strand displacement reaction as an example to show that the fluorophore alone can alter and even completely inhibit the dynamics. Strand-displacement is one of the best understood and highly specific methods of actuating large DNA devices, but still has a large potential for improvement with respect to kinetics. Addressing this challenge, we find that fluorophores tend to stabilize the equilibrium states of the system with different effects on its dynamics, whereby Cy-dyes are more prone to inhibit dynamics compared to Atto-dyes. Beyond fluorophore labelling, DNA nanotechnology uses a myriad of other chemical modifications on the DNA, form reactive amine or thiol groups, hydrophobic tags, spacers, photocleavable groups or modifications for click chemistry.^[47]^ We anticipate that our observations are not limited to dye molecules - these other chemical modification would very likely have similar effects. It is thus generally possible to shape the energy landscape for dynamic reconfiguration as well as the equilibrium configuration without changing the DNA sequence.

Our results are directly relevant for various applications that capitalize on dynamic DNA systems, from bottom-up synthetic biology to biosensing and the the increasingly popular superresolution technique DNA-PAINT.^[23]^ Without doubt, the possibility to precisely shape energy landscapes for dynamic DNA nanostructures will lead to metastable DNA nanostructures and fully reversible DNA devices with unprecedented complexity – mimicking the intricate workings of natural nanomachines.

## Acknowledgement

The authors acknowledge Tobias Abele for help with the analysis of the droplet’s peripheral intensity. K.G. acknowledges funding from the Deutsche Forschungsgemeinschaft (DFG, German Research Foundation) under Germany’s Excellence Strategy via the Excellence Cluster 3D Matter Made to Order (EXC-2082/1 - 390761711). The work was further supported by the DFG via the principal investigator grant IG 109/1-1 awarded to M.I. Computational resources were provided by the Max Planck Computing and Data Facility and the Leibniz Supercomputing Centre (Garching, Germany). K.J. thanks the Carl Zeiss Foundation for financial support. All authors acknowledge the Max Planck Society for its general support.

