## Supplementary Information for "Choice of fluorophore affects dynamic DNA nanostructures"

**References** **14**

### Supplementary Figures

#### Supplementary Figure S1: ssDNA/tsDNA-binding is equilibrated within the first hour

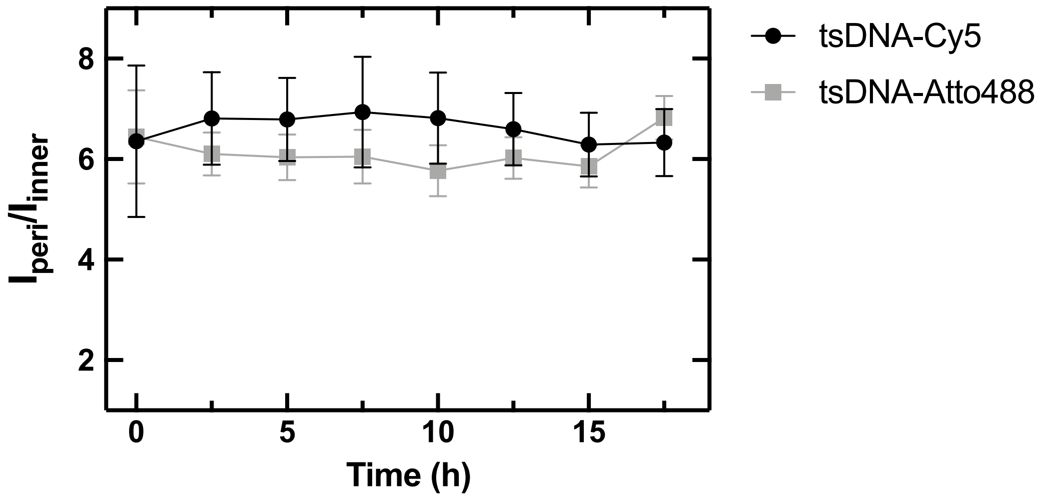

Figure 1: ssDNA/tsDNA-binding efficiency remains constant over time. Fluorescence intensity ratio $I_{peri}/I_{in}$ of tsDNA inside water-in-oil droplets for ssDNA-Cy3/tsDNA-Atto488 (gray) and ssDNA-Alexa488/tsDNA-Cy5 (black) over the course of 17.5 h. The fluorescence intensity ratios remain constant over time. This shows that the ssDNA/tsDNA binding reaches a dynamic steady-state already minutes after the droplet production and importantly before the imaging. Therefore, all experiments in this manuscript show the equilibrated state (with the exception of Figure 3, where we induce a pH change). Error bars correspond to the standard deviation of n=9 droplets.

#### Supplementary Figure S2: Fluorophore-tagged ssDNA does not interact with droplet-stabilizing surfactants

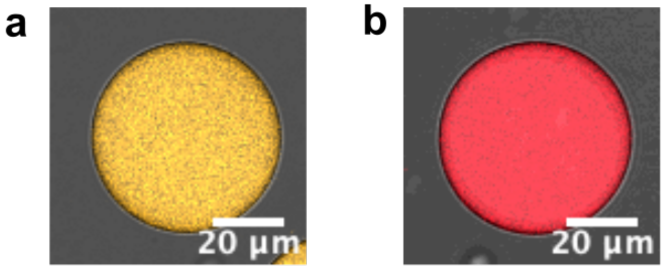

Figure 2: Fluorophore-tagged ssDNA does not interact with droplet-stabilizing surfactants. Representative confocal images of water-in-oil droplets containing Cy3-(**a**, $\lambda_{ex} = 561\,\mathrm{nm}$) and Cy5-labeled ssDNA (**b**, $\lambda_{ex} = 647\,\mathrm{nm}$) without cholesterol-modification at pH8. The solution contained 20 mM potassium phosphate buffer, 10 mM $MgCl_2$ and 1.5 µM DNA. Scale bars: 20 µm.

#### Supplementary Figure S3: MD simulations suggest that Cy5-labeling can stabilize more compact ssDNA conformations

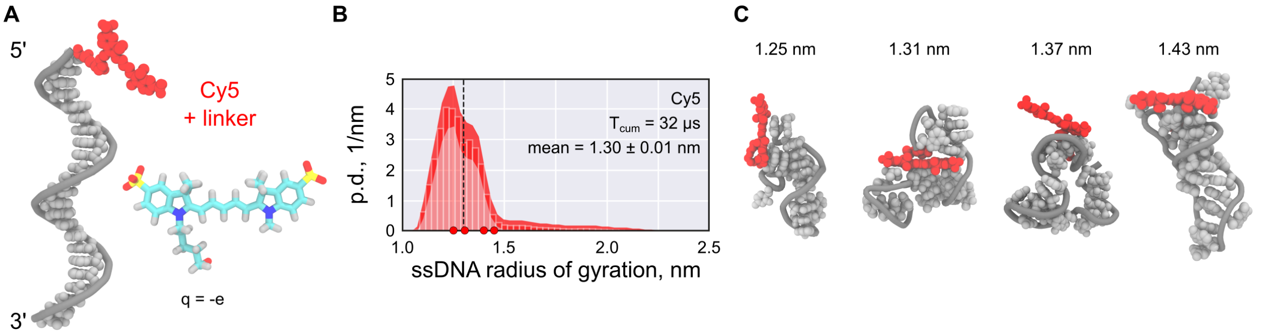

Figure 3: **A** Starting conformation of Cy5-(red) ssDNA (gray). The chemical structure of the fluorophore and its net charge are shown as an inset. **B** Probability density distributions of the gyration radius of ssDNA labeled with Cy5. The shaded area indicates the 95% confidence interval estimated using bootstrapping (see Methods in the main text). The black dashed line indicates the mean of the distribution, $T_{\mathrm{cum}}$ the cumulative simulation time. **C** Representative structure snapshots of the Cy5-ssDNA. Positions of the selected snapshots within the corresponding distributions are marked with dots in the probability density distribution.

#### Supplementary Figure S4: pH hysteresis of ssDNA/tsDNA binding

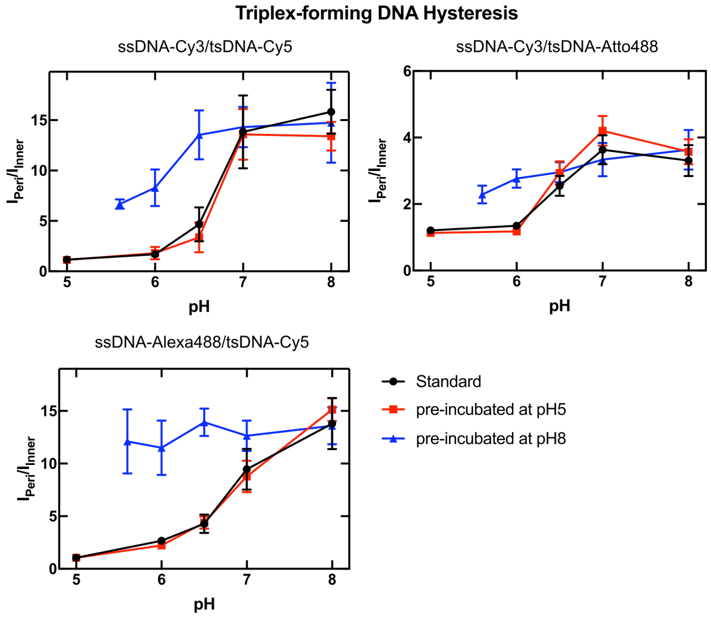

Figure 4: pH hysteresis of ssDNA/tsDNA binding. Fluorescence intensity ratio $I_{Peri}/I_{Inner}$ of tsDNA within water-in-oil droplets at different pH values for different ssDNA/tsDNA combinations including ssDNA-Cy3/tsDNA-Cy5, ssDNA-Cy3/tsDNA-Atto488 and ssDNA-Alexa488/tsDNA-Cy5. The tsDNA was incubated with ssDNA for 10 min in 10 mM sodium phosphate buffer and 20 mM $MgCl_2$ at pH 5 (blue curve), pH 8 (red curve) or directly at the indicated pH (black). After incubation the solutions were mixed 1:1 with 200 mM phosphate buffers ranging from pH 5 to 8 and subsequently encapsulated into droplets. The droplets were sealed in an observation chamber and imaged with confocal fluorescence microscopy. From these images, the fluorescence intensity ratio $I_{Peri}/I_{Inner}$ was extracted. The plots indicate that the duplex dissociation happens at lower pH values compared to the duplex formation. Importantly, this is independent of the type of fluorophore modification as it is visible for all tested ssDNA/tsDNA combinations. In some cases, the attachment to the periphery seems mostly irreversible (see ssDNA-Alexa488/tsDNA-Cy5). Note that it was not possible to revert back to pH 5 because the buffer range of the phosphate buffer used lies between pH 5.8 and pH 8.2. Error bars correspond to the standard deviation of n$\geq$25 droplets.

**Supplementary Figure S5: Light-mediated acidification of water-in-oil droplets reveals different detachment kinetics of tsDNA modified with different fluorophores.**

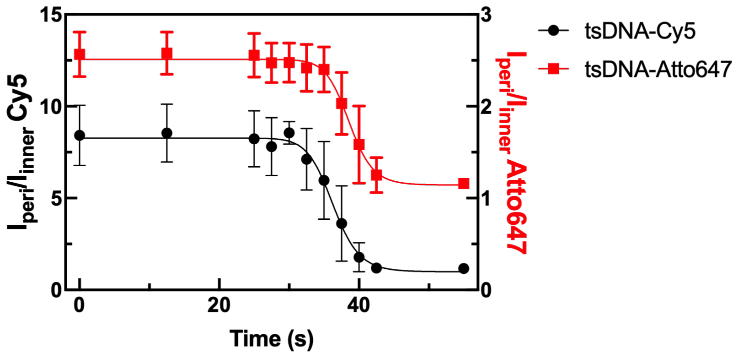

Figure 5: Light-mediated acidification of water-in-oil droplets reveals different detachment kinetics of tsDNA modified with different fluorophores. Fluorescence intensity ratio $I_{peri}/I_{inner}$ of tsDNA-Cy5 (black) and tsDNA-Atto647 (red) within droplets over time with Cy3-labeled ssDNA. Error bars correspond to the standard deviation of n $\geq$ 20 droplets for the DNA experiments. Solid lines represent sigmoidal fits with turning points at 36.3 $\pm$ 0.3 s (tsDNA-Cy5) and 38.6 $\pm$ 0.3 s (tsDNA-Atto647). tsDNA-Cy5 detaches faster from the droplet periphery during the acidification indicating that the fluorophore modification on tsDNA alone, changes the reaction kinetics.

### Supplementary Texts

#### Supplementary Text S1: Estimation of the apparent thickness of the reaction layer $\varepsilon$

The thickness of the reaction layer, $\varepsilon$, was determined by analyzing the intensity at the droplet periphery for the ssDNA/tsDNA-Cy5 combination at pH8. The intensity at the droplet periphery $I_{peri}$ was quantified by fitting a Gaussian distribution along a line orthogonal to the droplet circumference. The fit function was of the form $y = a + (b - a)e^{-(x-c)/2d^2}$, where $a$ is an offset parameter, $b$ is the maximum intensity, $c$ is the center point, and $d$ is the width. This analysis was repeated 20 times every 18° along the droplet circumference for 12 representative droplets. $\varepsilon$ was defined as the average of the full width at half maximum (FWHM) and calculated via $\varepsilon = 2\sqrt{2ln(2)}\,\bar{d}$ to be $\varepsilon = 1.82 \pm 0.37\,\mu$m.

#### Supplementary Text S2: Reaction-diffusion model of ssDNA-tsDNA interaction in a spherical volume

To describe the reaction-diffusion of the ssDNA and tsDNA molecules in a spherical droplet with a radius $r_0$, we adapted a previously proposed, volumetric model,[1,2] in which the binding and dissociation reactions are localized in a layer near the spherical droplet periphery. Outside of this layer, tsDNA is assumed to diffuse freely and isotropically. Let $\varepsilon$ denote the thickness of the reaction volume $V_\varepsilon \approx 4\pi r_0^2 \varepsilon$, in which ssDNA is located, $T(r,t)$ and $S(r,t)$ be the volumetric concentrations of *free* tsDNA and ssDNA, respectively, and $T_{tot}$ and $S_{tot}$ be the *total* concentrations of tsDNA and ssDNA in the droplet volume. Then, the equations describing the evolution of $T(r,t)$ and $S(r,t)$ take the form:

$$\begin{aligned} \frac{\partial T}{\partial t} &= D\Delta T - k_{on}TS + k_{off}(S_{tot} - S), \\ \frac{\partial S}{\partial t} &= -k_{on}TS + k_{off}(S_{tot} - S), \\ 0 \le r &\le r_0, t > 0, \end{aligned} \tag{1}$$

where $D$ is the diffusion constant of tsDNA, $k_{on}$ and $k_{off}$ are the association and dissociation rate constants, respectively. Since all binding sites (ssDNA) are concentrated within $V_\varepsilon$, the concentration of ssDNA must vanish outside of this area, *i.e.* $S(r,t) = 0$ for $r < r_0 - \varepsilon$. Furthermore, we assume that the droplet boundary is impermeable, *i.e.* $\partial T/\partial r|_{r_0} = 0$. We also note that, strictly speaking, Eq. 1 is solved separately for $0 \le r < r_0 - \varepsilon$ and $r \ge r_0 - \varepsilon$, but the joint solution must be continuous and differentiable everywhere on $0 \le r \le r_0$. We therefore require that:

$$\begin{aligned} T^{(0)}|_{r_0-\varepsilon} &= T^{(\varepsilon)}|_{r_0-\varepsilon}, \\ \frac{\partial T^{(0)}}{\partial r}|_{r_0-\varepsilon} &= \frac{\partial T^{(\varepsilon)}}{\partial r}|_{r_0-\varepsilon}, \end{aligned} \tag{2}$$

where $T^{(0)}$ and $T^{(\varepsilon)}$ are the piecewise solutions of $T(r,t)$ within and outside of $V_\varepsilon$, respectively.

We seek for the steady-state solution of Eq. 1 because it reflects experiments in which the ssDNA-tsDNA binding has attained equilibrium (see, *e.g.*, Figure 1 in the main text and Figure S1). The steady-state solution is given by $T(r,t) = T_{eq}(r)$ and $S(r,t) = S_{eq}(r)$. Solving Eq. 1 under Eq. 2 and the boundary conditions yields:

$$\begin{aligned} T_{eq}(r) &= C, \\ S_{eq}(r) &= \frac{K_D S_{tot}}{T_{eq}(r) + K_D}, \end{aligned} \tag{3}$$

where we have introduced the apparent dissociation constant $K_D = k_{off}/k_{on}$, and where $C$ is an arbitrary constant. This constant can be found explicitly using the mass conservation of tsDNA within the droplet volume, that is:

$$\underbrace{\int_0^{r_0} T_{eq}(r)4\pi r^2 dr}_{\text{free tsDNA}} + \underbrace{\int_0^{r_0} (S_{tot} - S_{eq}(r))4\pi r^2 dr}_{\text{bound tsDNA}} = \underbrace{T_{tot}\frac{4}{3}\pi r_0^3}_{\text{total tsDNA}}. \tag{4}$$

By substituting Eq. 3 into Eq. 4 and assuming that $\varepsilon \ll r_0$, we finally obtain the explicit expressions for the steady-state concentrations:

$$\begin{aligned} T_{eq} &= -\frac{1}{2}\left[3\frac{\varepsilon}{r_0}S_{tot} + K_D - T_{tot}\right] + \sqrt{\left[3\frac{\varepsilon}{r_0}S_{tot} + K_D - T_{tot}\right]^2 + 4T_{tot}K_D}, \\ S_{eq} &= \frac{K_D S_{tot}}{T_{eq} + K_D}. \end{aligned} \tag{5}$$

In the experiment, the concentrations $T_{eq}$ and $S_{eq}$ are not available directly, but only through the respective fluorescent intensities measured in the droplet interior and at the periphery with confocal microscopy. More specifically, the intensity of labelled tsDNA molecules is $I = E_{ex}E_{em}V_{obs}T_{eq}$,[3,4] where $V_{obs}$ is the observation volume, and $E_{ex}$ and $E_{em}$ are excitation and emission functions of the light path, respectively. Assuming that imaging settings do not change for the tested fluorophore combinations, the ratio between the peripheral, $\tilde{I}_{peri}$, and inner intensity, $\tilde{I}_{in}$, of tsDNA fluorescence can be expressed as:

$$\frac{\tilde{I}_{peri}}{\tilde{I}_{in}} = \frac{(S_{tot} - S_{eq}) + T_{eq}}{T_{eq}} = 1 + \frac{S_{tot}}{T_{eq} + K_D} \equiv A(\varepsilon, K_D). \tag{6}$$

Here, $\tilde{I}_{peri} = I_{peri}/(2\pi r_0 \varepsilon h)$ and $\tilde{I}_{in} = I_{in}/(\pi r_{in}^2 h)$, where $h$ is the characteristic height of the point spread function of the microscope (typically a few microns) and $r_{in} < r_0$ is the radius of an arbitrary sphere placed within the droplet volume. The expression in Eq. 6 can be now employed to estimate $K_D$ by experimentally measuring $A$ and precomputing $\varepsilon$ (see Supplementary Text S2). By analogy, the uncertainty of $K_D$ can be calculated using error propagation on A:

$$\sigma_A^2 \approx \left(\frac{\partial A}{\partial \varepsilon}\right)^2 \sigma_\varepsilon^2 + \left(\frac{\partial A}{\partial K_D}\right)^2 \sigma_{K_D}^2, \tag{7}$$

where the partial derivatives are given by:

$$\begin{aligned} \frac{\partial A}{\partial \varepsilon} &= \frac{3S_{tot}^2}{2r_0(T_{eq}+K_D)^2}\left[1 - \frac{3\frac{\varepsilon}{r_0}S_{tot}+K_D-T_{tot}}{\sqrt{(3\frac{\varepsilon}{r_0}S_{tot}+K_D-T_{tot})^2+4T_{tot}K_D}}\right], \\ \frac{\partial A}{\partial K_D} &= -\frac{S_{tot}}{2(T_{eq}+K_D)^2}\left[1 + \frac{3\frac{\varepsilon}{r_0}S_{tot}+K_D+T_{tot}}{\sqrt{(3\frac{\varepsilon}{r_0}S_{tot}+K_D-T_{tot})^2+4T_{tot}K_D}}\right]. \end{aligned} \tag{8}$$

### Supplementary Videos

#### Supplementary Video S1: MD trajectory of unlabelled ssDNA

MD trajectory showing the evolution of the starting conformation of the unlabelled ssDNA over the total time of 5 $\mu$s and sampled every 5 ns. The conformation of the DNA remains, on average, more extended throughout the simulation than the labeled counterparts, thus being more accessible for complementary base pairing. See Fig. 2B for the distribution of the radius of gyration.

#### Supplementary Video S2: MD trajectory of Cy3-ssDNA

MD trajectory showing the evolution of the starting conformation of the Cy3-labelled ssDNA over the total time of 6 $\mu$s and sampled every 5 ns. The Cy3-ssDNA adopts more compact conformations compared to the unlabeled counterpart, in which the hydrophobic bases are wrapped around the Cy3 fluorophore, reducing their accessibility for complementary base pairing. See Fig. 2B for the distribution of the radius of gyration.

#### Supplementary Video S3: MD trajectory of Cy5-ssDNA

MD trajectory showing the evolution of the starting conformation of the Cy5-labelled ssDNA over the total time of 6 $\mu$s. Like in the case of the Cy3 modification, Video S2, the ssDNA adopts more compact conformations, where the bases are wrapped around Cy5 fluorophore, reducing their accessibility for complementary base pairing. See Fig. S3B for the distribution of the radius of gyration.

#### Supplementary Video S4: MD trajectory of Alexa488-ssDNA

MD trajectory showing the evolution of the starting conformation of the Alexa488-labelled ssDNA over the total time of 6 $\mu$s. The accessibly of the bases is, on averaged, higher compared to the Cy3 and Cy5 modifications (Video S2 and S3), possibly due to weaker dye-ssDNA stacking interactions and the higher negative charge of Alexa 488. See Fig. 2B for the distribution of the radius of gyration.

#### Supplementary Video S5: Light-mediated detachment kinetics of Cy5-tsDNA

Representative confocal time series of Cy5-labeled tsDNA ($\lambda_{ex} = 647\,\text{nm}$) encapsulated together with unlabelled ssDNA and NPE-caged-sulfate into water-in-oil droplets at pH 8. Upon illumination with 405 nm for 50 s, uncaging of NPE-caged-sulfate releases a proton, leading to acidification inside the droplet. During acidification, tsDNA-Cy5 detaches from the droplet periphery as the triplex state is energetically favoured at low pH. The normalized inner intensity $\mathrm{I_{in}}$ over time is shown in Figure 3**C** (Main). in Scale bars: 20 µm.
